# Temporal prediction captures key differences between spiking excitatory and inhibitory V1 neurons

**DOI:** 10.1101/2024.05.12.593763

**Authors:** Luke Taylor, Friedemann Zenke, Andrew J. King, Nicol S. Harper

**Author notes:** Contributing authors.

## Abstract

Neurons in primary visual cortex (V1) respond to natural scenes with a sparse and irregular spike code that is carefully balanced by an interplay between excitatory and inhibitory neurons. These neuron classes differ in their spike statistics, tuning preferences, connectivity statistics and temporal dynamics. To date, no single computational principle has been able to account for these properties. We developed a recurrently connected spiking network of excitatory and inhibitory units trained for efficient temporal prediction of natural movie clips. We found that the model exhibited simple and complex cell-like tuning, V1-like spike statistics, and, notably, also captured key differences between excitatory and inhibitory V1 neurons. This suggests that these properties collectively serve to facilitate efficient prediction of the sensory future.

## Introduction

Our visual system allows us to navigate the world and interpret the most unlikely of scenes, like a toy brachiosaurus sitting atop a brick wall. The primary visual cortex (V1) plays a central role in processing diverse and complex stimuli, with neurons tuned to the orientation and motion of moving edges and shapes [1–3]. These features are communicated by a sparse and irregular spike code [4] that is carefully balanced by interactions between the cortex’s excitatory and inhibitory neural subpopulations [5–8]. There is growing evidence for the idea that visual perception arises from the brain engaging in predictive processing [9]. In particular, recent research suggests that neurons in sensory brain areas, such as V1, may have evolved to represent features in natural stimuli that are predictive of the immediate sensory future [10, 11]. Such representations are evolutionary advantageous, since they allow inferences about the underlying causes of sensory signals, which may help to guide future action, and eliminate unnecessary sensory information [12].

Recent computational studies have demonstrated that feedforward artificial neural networks (ANNs) trained for temporal prediction using natural movies develop units with receptive field properties that resemble those of V1 neurons [10, 11]. Specifically, these units replicate the spatial tuning of V1 neurons and notably also their temporal dynamics - a property not readily captured by other models [10]. However, while feedforward ANNs loosely resemble V1’s biological circuit [13], they omit several key features, including the recurrent connectivity of the cortex; they do not conform to Dale’s law [14], where neurons are either excitatory or inhibitory; and they omit the brain’s key physiological unit of computation: the spike. It therefore remains unclear to what extent - if at all - a biologically detailed model of V1 optimized for temporal prediction would capture the physiological properties of its neurons.

To bridge this shortcoming, we developed a spiking model of recurrently-connected excitatory and inhibitory units trained on natural movies under metabolic-like constraints to predict the sensory future from recent spike activity. We found the units in our model to exhibit simple and complex cell-like tuning. Extending beyond prior results, the spike statistics of our model resembled those of V1 neural responses to natural stimuli. Notably, the excitatory and inhibitory units in our model captured the differences in spike statistics, tuning preferences, connectivity statistics and temporal dynamics between excitatory and inhibitory V1 neurons.

## Results

### The spiking V1 model

We implemented our model as a population of recurrently-connected spiking units and adopted the leaky integrate-and-fire (LIF) model to mimic real neural spiking dynamics [15] (Figure 1A). Similar to the ratio of excitatory-to-inhibitory neurons in V1 [5–8], we set 85% of these units as excitatory and the remaining 15% as inhibitory. The model’s input consists of a sequence of 20 *×* 20px patches obtained from retina-filtered natural movies recorded at 120 frames-per-second. At each time step, the model linearly translates a 126ms (15 frames) span of stimulus history into a distinct input current to each unit. The 126ms duration covers the potential integration span and spread of latencies of the incoming thalamocortical input [16]. The model was trained to linearly predict the spatial movie patch 42ms (5 frames) into the future using a 16ms (2 frames) span of past population spike activity (Figure 1B). We chose the 42ms prediction target to be of similar duration to the minimum latency of visual input to V1 [17–20] and of similar duration to the response latency of the fastest visuomotor actions [21].

**Fig. 1:**
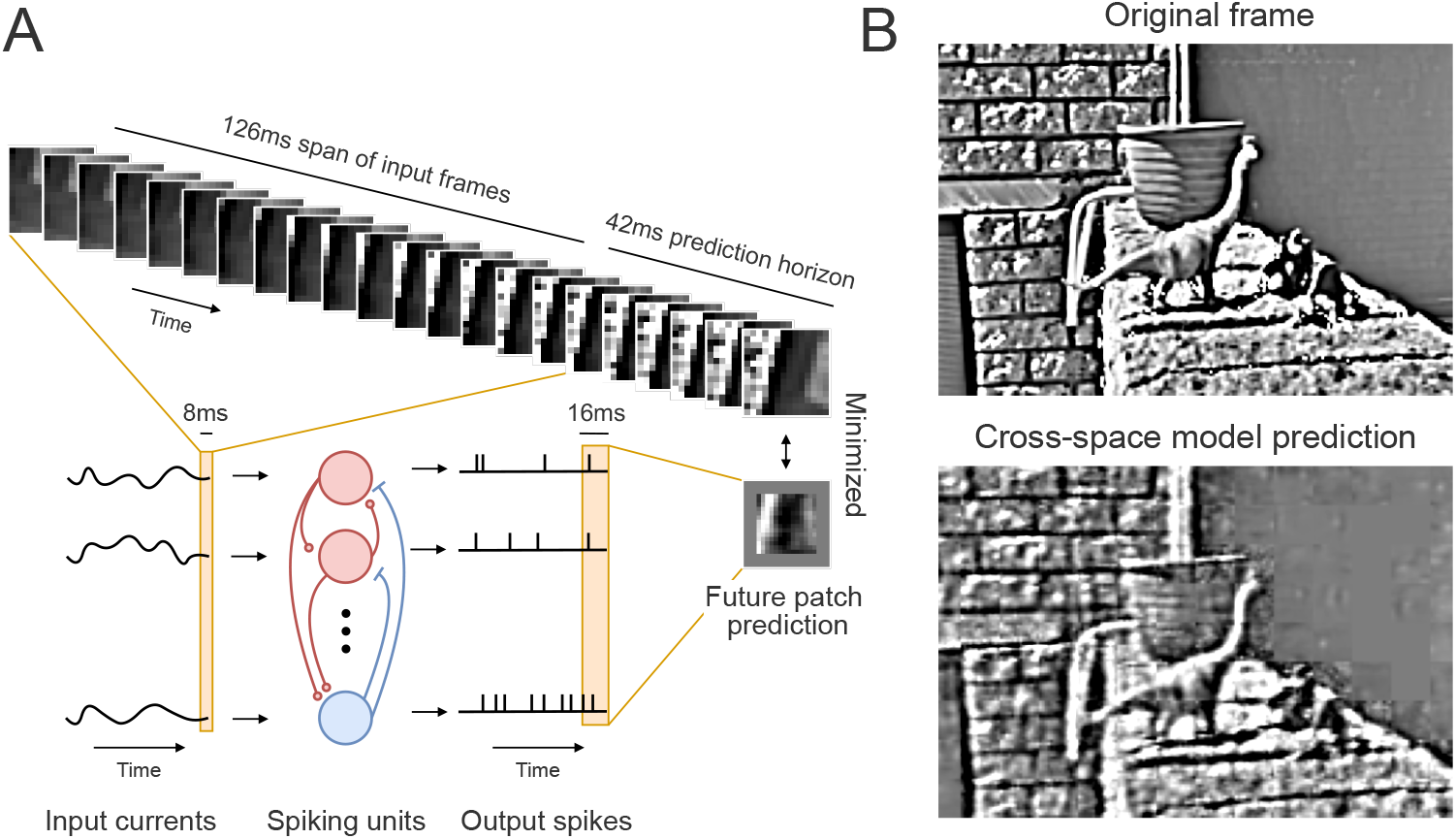
The spiking temporal prediction model with recurrently connected excitatory and inhibitory units. **A**. Schematic of our model, which at every time step linearly maps a 126ms span of natural movie frame patches of the past into a distinct input current value to every spiking unit. A 16ms span of population output spikes is linearly mapped to create a prediction of the spatial movie patch 42ms into the future. Red units are excitatory and blue units are inhibitory. **B**. An example movie frame of a toy dinosaur sitting on a brick wall, which was not included in the training data (top), with the model’s reconstruction using multiple patch predictions across different spatial locations (bottom).

To emulate the metabolic energy constraints of the brain, we also trained our model to reduce a metabolic-like cost whilst performing the frame prediction (see Methods). This approximated the activity-induced cost at the model weights, similarly to the energy expenditure of neural synapses. We made an assumption about the brain in our metabolic-like cost function, namely that inhibitory V1 neurons consume less energy than their excitatory counterparts per spike. We found this assumption to be important for obtaining all of the V1-like tuning differences between the excitatory and inhibitory model units. Lastly, training was performed using a variant of the backpropogation algorithm [22], known as surrogate gradient learning [23], which permits gradient descent in non-differentiable spiking networks. This optimization process determined the model’s connectivity and the membrane time constant of each spiking unit. We remain agnostic about the biological plausibility of this training method, and how cortex might optimize for the objective of temporal prediction.

### Emergence of a V1-like spike code

We examined the model’s spike activity in response to retina-filtered clips taken from the Star Wars movies that were not used for training, and found a number of V1-like response characteristics to emerge. We observed the model’s spike responses to be sparse and diverse, with variability in the number and timing of spikes across units (Figure 2A). Further quantifying the spike responses revealed a firing rate distribution with an exponential fall-off (Figure 2B), as has similarly been reported in monkey [4] and cat V1 [24]. The average model’s firing rate was also comparable to mean V1 firing rates observed across different animal species in response to natural stimuli (model 6.57 *±* 14.68Hz; anesthetized monkey V1: 5.06 *±* 0.75Hz [4]; anesthetized cat V1: 3.96 *±* 3.61Hz [24] and awake cat V1:8.9 *±*7Hz [25]). Additionally, the inhibitory model units exhibited a higher mean firing rate compared to the excitatory model units (Figure 2C). Such differences in firing rates between excitatory and inhibitory neurons to visual stimulation have also been reported in mouse V1 [26].

**Fig. 2:**
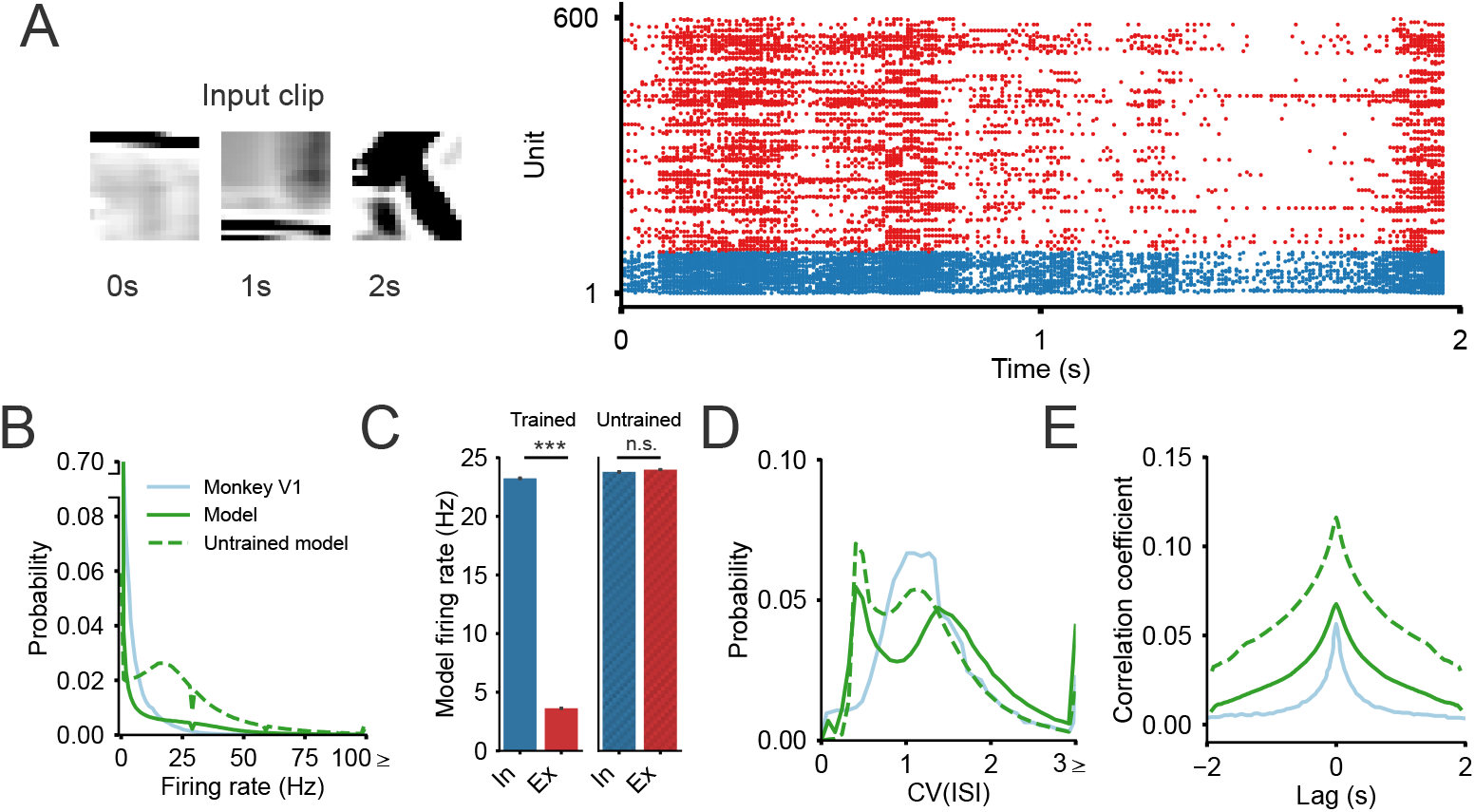
Spiking statistics of the model resemble those of V1. **A**. Input movie clip (left) and resulting model spike raster (right). Red and blue dots correspond to the excitatory and inhibitory units, respectively. **B**. Firing rate distribution of the trained and untrained model units and monkey V1 neurons. **C**. Difference in the model’s firing rates between the inhibitory (In) and excitatory (Ex) units in the trained (left) and untrained model (right). Bars plot the mean and standard error over units and input clips, with statistical significance assessed using the Mann-Whitney U test (\*\*\**p <* 0.001). **D**. Coefficient of variation of the interspike intervals CV(ISI) distribution of the trained and untrained model units and monkey V1 neurons. **E**. Cross-correlogram of the trained and untrained model units and monkey V1 neurons. Monkey V1 data in all plots were obtained from [4] in response to monkey’s watching segments of the Star Wars movies.

The model’s spike responses were highly variable, exhibiting a low degree of correlation between the units. We examined the variability in the spike responses using the coefficient of variation of the interspike intervals CV(ISI) (Figure 2D), which is defined as the ratio of the ISI standard deviation to the ISI mean. A spike train with a CV(ISI) below unity is more regular than a spike train with a CV(ISI) above unity. We observed a large range of variability in the spike trains, with sampled spike trains exhibiting more irregular (65% with CV(ISI) ≥ 1) than regular (35% with CV(ISI)*<* 1) firing patterns, reminiscent of the spike variability observed in cat [27] and monkey V1 [4, 28].

We quantified the correlation between the different unit spike trains by calculating the spike train cross-correlogram, which measures the correlation between randomly-sampled unit pairs at different time lags (Figure 2E). We found the model units to have a low peak correlation (*CC* = 0.07 at lag 0s), with the correlation symmetrically decaying for increasing and decreasing lag. Similar correlation statistics have also been reported in anesthetized monkey [4] and in awake mouse V1 [29]. Lastly, the resemblance of the model’s spiking statistics to the biology appeared to be a direct consequence of optimizing the model for efficient temporal prediction, as evident by the untrained model’s firing rates (Figure 2B and C) and spike correlations (Figure 2E) being less similar to V1 than the trained model. However, we did notice a slight divergence in the trained model’s spike variability compared to V1 (Figure 2D).

### Emergence of simple and complex cell-like tuning

We obtained model unit receptive fields (RFs) using a spike-triggered average to spatiotemporal white-noise input clips, and observed a considerable number of similarities between the model unit RFs and the RFs of V1 neurons across different animal species. Qualitatively, we found many units that resemble simple cells [1, 3], with stereotyped RFs consisting of varying number of excitatory and inhibitory subfields tuned to a particular orientation and spatial frequency [32, 33] (Figure 3A). The temporal structure of these RFs also resemble those of V1 neurons, as evident in their decay into the past (Figure 3B); their polarity profile, with RF polarity either changing (Figure 3B units 1, 4 and 5) or remaining fixed over time (Figure 3B units 2 and 3) [35–37]; and their spatiotemporal structure, with RFs either being space-time separable (Figure 3B units 1, 2 and 3) or space-time inseparable (Figure 3B units 4 and 5) [35].

**Fig. 3:**
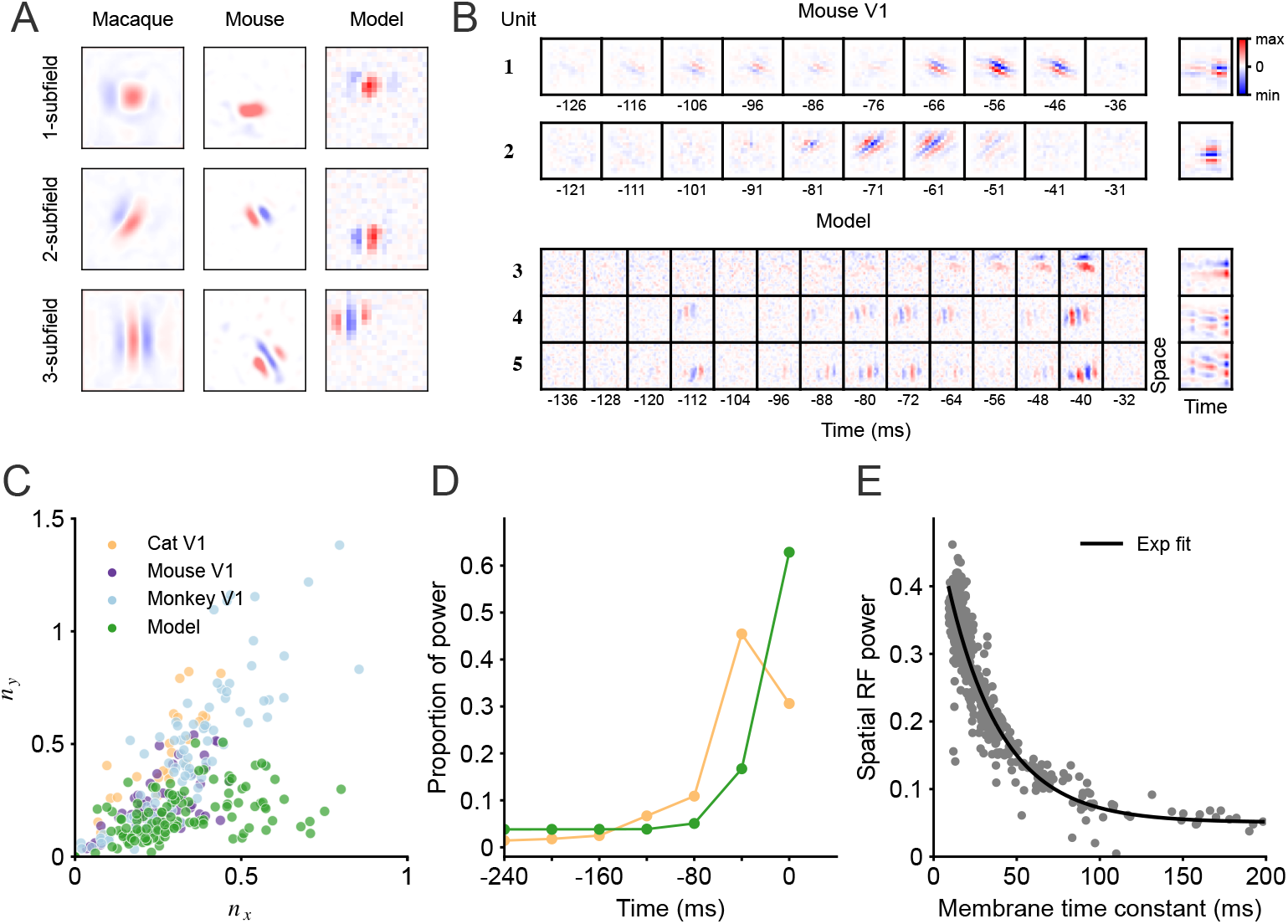
Spatial and temporal receptive field properties of the model resemble V1. **A**. Example spatial RFs with varying number of subfields of macaque V1 neurons [30], mouse V1 neurons [26] and model units. **B**. 1 − 2: example space-time separable spatiotemporal RFs of mouse V1 neurons [31]. 3 − 5: example spatiotemporal RFs of model units (3: space-time separable and 4 − 5: space-time inseparable). Right column: corresponding spatiotemporal RFs obtained by summing along the RFs’ axis of orientation (see Methods). **C**. Distribution of spatial RF shapes from V1 neurons in cats [32], mice [26], monkeys [33] and model units. Each model unit’s spatial RF was taken at its time point of highest power. The x-axis (*n*_*x*_) and y-axis (*n*_*y*_) are a measure proportional to the number of RF subfields and their respective lengths [33]. **D**. Temporal RF power profile of model units and V1 cat neurons [34], quantified as the sum of squared weights over space and averaged across the population over ∼ 40ms time bins. **E**. Scatter plot of the spatial RF power (largest mean squared values of a spatial RF in time) and membrane time constant of the model units. Dark line is the fitted negative exponential curve (*R*^2^ = 0.85).

We fit each model unit’s spatial RF at the time with the highest power using a Gabor function to further quantify its shape, orientation and spatial frequency. The distribution over RF shape characteristics, as defined by the width *n*_*x*_ and height *n*_*y*_ of the RF relative to its spatial frequency, matches the neural data (Figure 3C). We found the RF shapes to extend along a 1D-curve, from blob-like RFs at the origin (Figure 3A first row) to elongated bars with multiple subfields that are located further away from the origin (Figure 3A last row). Our model captures a large portion of the blob-like RFs, which constitute a substantial part of the neural data (with *n*_*x*_ and *n*_*y*_ *<*∼ 0.25). It also captures units with multiple subfields (*n*_*x*_ *>* ∼ 0.25), although the subfields are typically less elongated than in the biology. We found the model RFs to be most similar to the mouse V1 RFs, as measured by the Euclidean distance between the model RF-shape centroid and the different animal RF-shape centroids (cat [32] centroid (0.26, 0.46) with distance 0.31; mouse [26] centroid (0.25, 0.24) with distance 0.09; monkey [33] centroid (0.33, 0.43) with distance 0.29; model centroid (0.26, 0.15)). The temporal profile across all model RFs also exhibited clear similarities to the neural data, as characterized by a decaying power profile (mean squared RF values over space, units and ∼ 40ms time bins) (Figure 3D).

We made an additional observation in our model, for which we found no prior reports in V1, by identifying a negative exponential relationship between the maximum spatial RF power and the membrane time constant of a unit (Figure 3E). The membrane time constant determines the operational timescale of a neuron, where neurons with smaller membrane time constants operate on shorter timescales than neurons with larger membrane time constants. Thus, translating our finding to the biology suggests that V1 neurons with a greater sensitivity to visual input operate on shorter timescales than neurons that exhibit a weaker stimulus sensitivity.

In addition to simple cells, V1 is also distinguished by another significant category of cells known as complex cells [2]. Like simple cells, complex cells are tuned for oriented edges and gratings. Unlike simple cells, however, complex cells are spatially invariant and respond to an oriented pattern regardless of its precise location in the neuron’s RF. Using full-field drifting sinusoidal gratings, we qualitatively identified units with response characteristics reminiscent of both simple and complex cells. We observed unit responses to oscillate over time to the movement of their optimal grating. Like simple cells, some model units displayed a high phase dependence with pronounced variations in their response to grating movement [38] (Figure 4A). In contrast, other units exhibited a larger mean response over time and were less influenced by the grating phase, consistent with the description of complex cells [39] (Figure 4B).

**Fig. 4:**
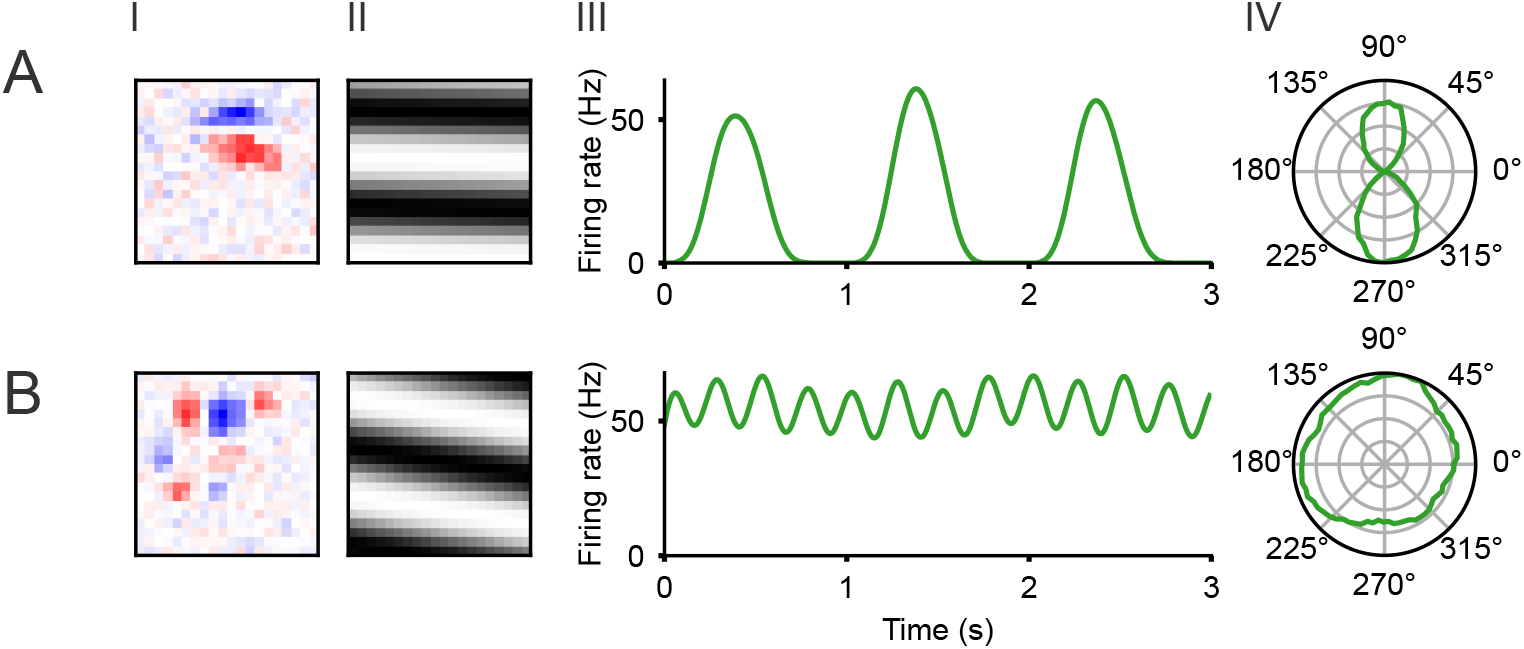
Units resembling simple and complex cells are found in the model. **A**. Example simple cell-like unit, including its RF (I); optimal grating (II); response to its optimal grating (III); and the average response of the unit over time plotted in relation to grating orientation (in degrees) for gratings presented at the unit’s preferred spatial and temporal frequency (IV). The optimal grating is defined as the full-field moving grating with an orientation and spatial and temporal frequency that give the highest average spike rate. **B**. Example complex cell-like unit.

We further quantified the distribution of simple and complex cell-like units in our model by calculating their modulation ratio *F*_1_*/F*_0_. This ratio is defined as the quotient between the amplitude of the first harmonic (*F*_1_) and the mean value (*F*_0_) of a unit’s response to its optimal grating, with a value below one classifying a response as complex cell-like [40, 41]. The distribution of modulation ratios in our model is bimodal (Figure 5A), which has similarly been reported in V1 neurons of monkeys [42] and mice [26, 43]. Intriguingly, we found the inhibitory units (median *F*_1_*/F*_0_ = 0.20) to exhibit more complex cell-like responses compared to the excitatory units (median *F*_1_*/F*_0_ = 1.62) (*p* = 3.59 *×* 10^−37^; Mann-Whitney U test), a difference also found between inhibitory and excitatory V1 neurons in mice [26].

**Fig. 5:**
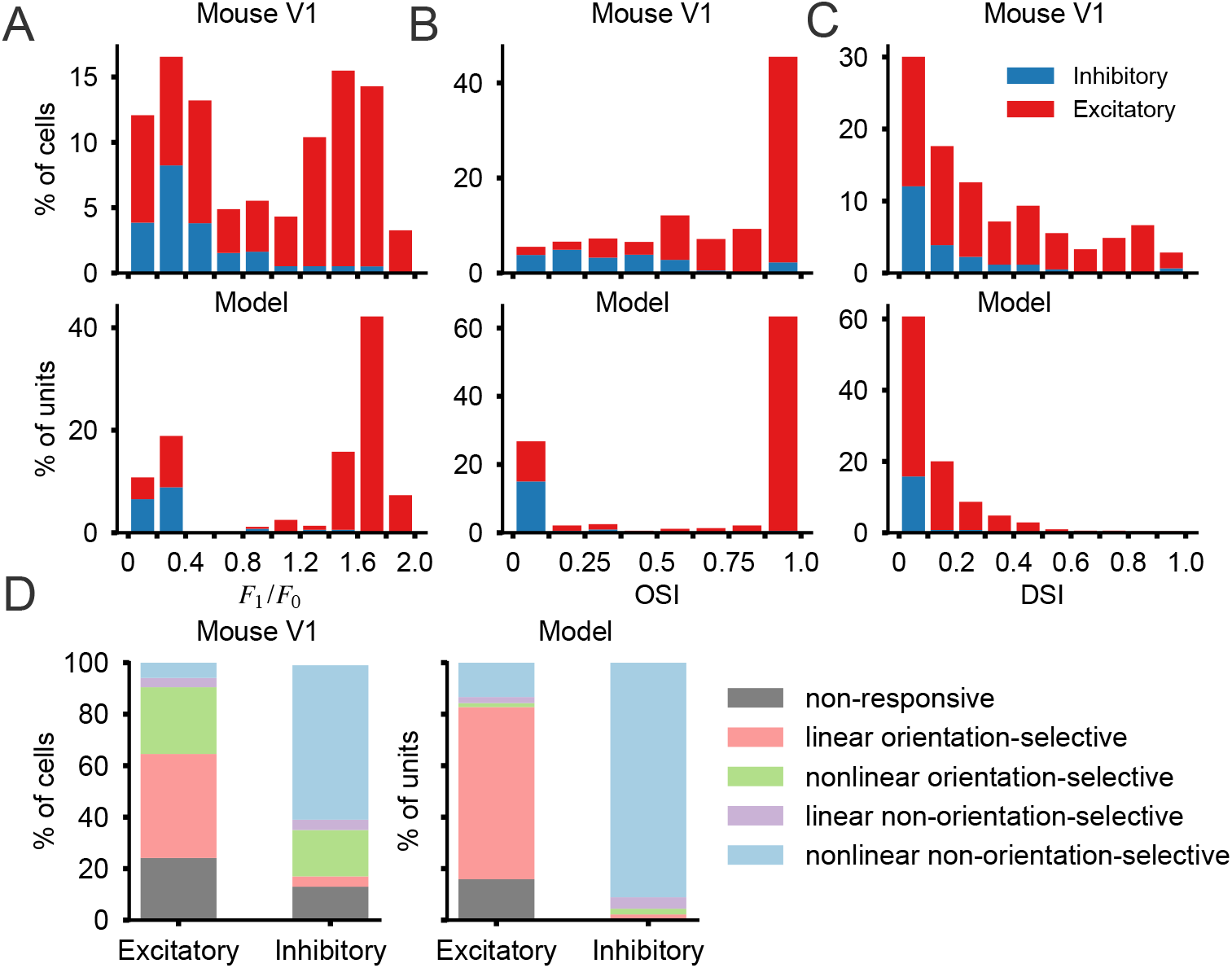
Tuning measures of the model bear resemblance to those in V1. **A**. Distribution of modulation ratios *F*_1_*/F*_0_. **B**. Distribution of orientation selectivity. **C**. Distribution of direction selectivity. **A-C** showing tuning properties of mouse V1 neurons [26] (top) and model units (bottom) to drifting gratings. **D**. Distribution of functional response categories for excitatory and inhibitory neurons in mouse V1 [26] (left) and for excitatory and inhibitory model units (right).

The model units exhibited similar orientation and direction preferences to those observed in V1 neurons in mice, as measured by the orientation selectivity index (OSI) and direction selectivity index (DSI), respectively [26, 44] (see Methods). Both measures range between zero and one, with a value of one corresponding to maximum selectivity. We found most units to have a strong preference for particular orientations (*OSI >* 0.5: 68%), with many units exhibiting near complete selectivity, as has similarly been reported in mouse V1 neurons (*OSI >* 0.5: 74%) [26] (Figure 5B). Capturing another difference between inhibitory and excitatory mouse V1 neurons [26], we found the inhibitory units (median *OSI* = 0.03) to be more broadly tuned than the excitatory units (median *OSI* = 1.00) (*p* = 2.14 *×* 10^−43^; Mann-Whitney U test). Additionally, we found the model units to have limited selectively for direction (*DSI <* 0.5: 97%), as has similarly been reported for mouse V1 neurons (*DSI <* 0.5: 77%) [26] (Figure 5C).

We classified units into different functional response categories, classifying a unit as linear (*F*_1_*/F*_0_ ≥ 1) or non-linear (*F*_1_*/F*_0_ *<* 1), and as orientation selective (*OSI* ≥ 0.5) or non-orientation selective (*OSI <* 0.5) using the modulation ratio and OSI, respectively. The largest functional response categories in the model for both the inhibitory and excitatory units were the same as the largest functional response categories for inhibitory and excitatory neurons in mouse V1 [26] (Figure 5D). Specifically, most inhibitory model units are non-linear and non-orientation selective (91%), as are most inhibitory neurons in mouse V1 (60%), while most excitatory model units are linear and orientation selective (66%), as are most excitatory neurons in mouse V1 (40%).

### Mirroring V1-cell physiology: membrane dynamics, connectivity and EI balance

V1 is characterized by distinct physiological differences between inhibitory and excitatory neurons, such as the timescales they operate at [45] and their connectivity statistics [8]. We found our model units to mirror these key differences between the inhibitory and excitatory neurons. Inhibitory neurons tend to operate at shorter timescales than excitatory neurons, as captured by differences in their membrane time constants. Mouse V1 inhibitory neurons exhibit a median membrane time constant of 9.76ms, whilst excitatory mouse V1 neurons have a notably longer median membrane time constant of 21.10ms [45] (*p* = 3.43 *×* 10^−47^; Mann-Whitney U test). Similar to mouse V1, we found the inhibitory and excitatory model units to operate at different timescales, with the inhibitory and excitatory units having a median membrane time constant of 12.64ms and 22.33ms, respectively (*p* = 1.46 *×* 10^−30^; Mann–Whitney U test) (Figure 6A).

**Fig. 6:**
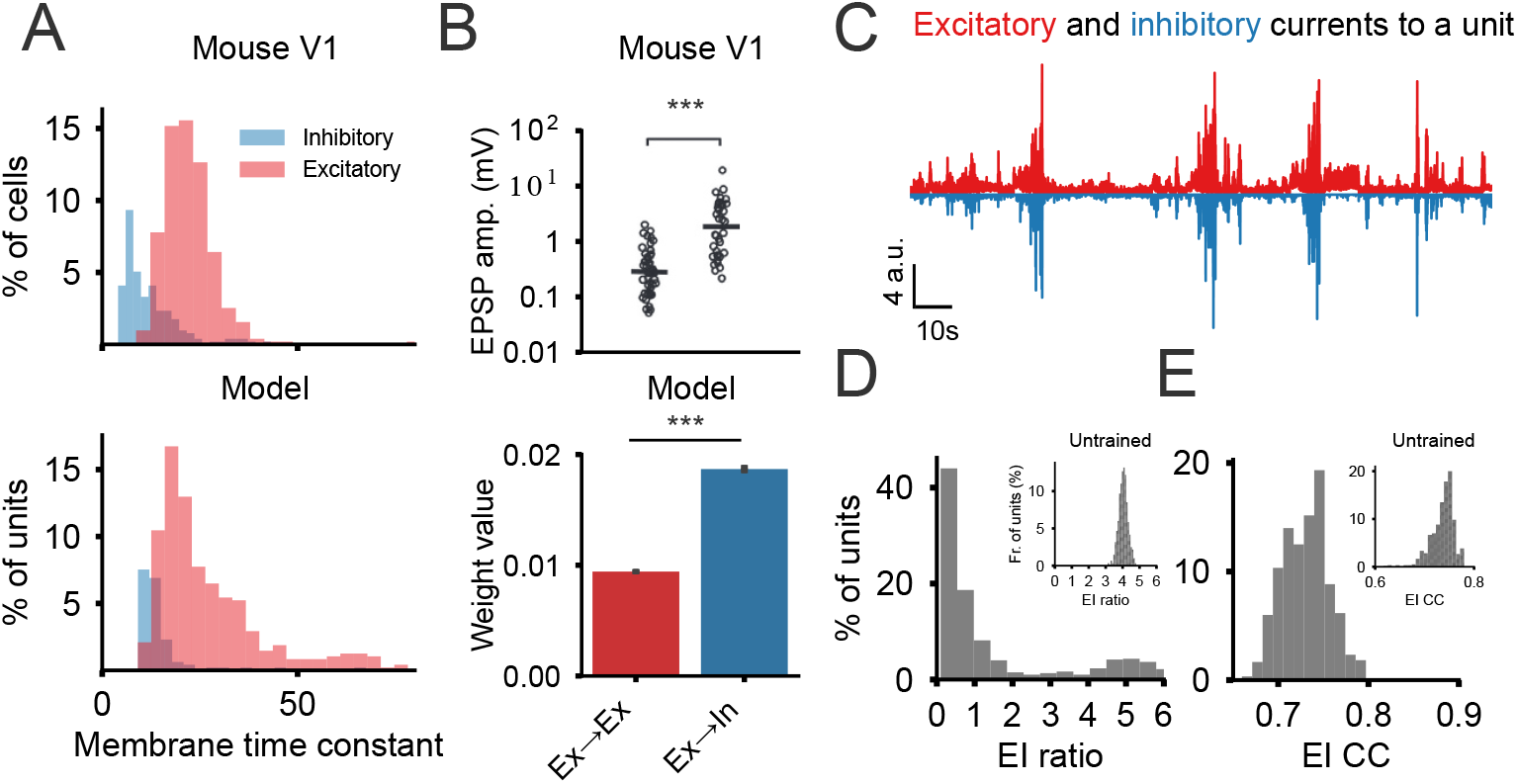
Inhibitory and excitatory model units operate at different temporal scales, with distinct connectivity statistics and balanced excitatory and inhibitory currents, like in V1. **A**. Membrane time constant distribution of inhibitory and excitatory neurons in mouse V1 (layer 4) [45] (top), and of inhibitory and excitatory model units (bottom). Membrane time constants of 122 excitatory model units with a value above 80ms are not shown. **B**. Top: EPSP amplitudes between excitatory neurons (Ex → Ex) and from excitatory neurons to inhibitory neurons (Ex → In) in mouse V1 (black lines plot median amplitudes) [8]. Bottom: model weight values between excitatory units and from excitatory to inhibitory units. Bars plot the median and bootstrapped standard error, with statistical significance assessed using the Mann-Whitney U test (\*\*\**p <* 0.001). **C**. Net-excitatory (top) and net-inhibitory (bottom) currents of an example model unit whilst inputting natural movie clips into the model. **D**. Distribution of global EI balance in the model. **E**. Distribution of precise EI balance in the model. Insets in **D**. and **E**. showcase the EI balances in the untrained model.

Excitatory V1 neurons have been shown to synapse more strongly onto inhibitory V1 neurons than other excitatory neurons in mice [8] and rats [46], as measured by their excitatory postsynaptic potential (EPSP) amplitude. We made a similar observation in our model, with the excitatory units connecting more strongly with the inhibitory units than with other excitatory units, as measured by their absolute connection weight (*p* = 0; Mann-Whitney U test) (Figure 6B).

A fundamental aspect in neurophysiology is the concept of cortical neurons receiving a balance of excitatory and inhibitory (EI) synaptic currents, as reported in somatosensory cortex [47], primary auditory cortex [48] and V1 [49–51]. This equilibrium is crucial for ensuring stability in neural activity [52] and proper cortical function [53]. We qualitatively observed a balance between the excitatory and inhibitory currents in the model units to held-out natural input movie clips (Figure 6C). EI balance can be quantified as global when the average excitatory and inhibitory currents are equal over time, and as precise, when the excitatory and inhibitory currents are coupled in time, in which each excitatory input to a neuron arrives simultaneously with an inhibitory counterpart [54]. We found the model units to exhibit a median global EI ratio of 0.67 over the held-out natural input movie clips (Figure 6D), where an EI ratio below unity has similarly been reported in mouse V1 [51, 55, 56]. Furthermore, we found the model units to exhibit a precise EI balance, as indicated by a notable correlation between the net-excitatory and net-inhibitory currents over clips and time across the model units (median: 0.73) (Figure 6E). Lastly, we found the excitatory and inhibitory currents in the model became more balanced during training for efficient temporal prediction, as evident in the untrained model having a notably larger median EI ratio of 4.04 (Figure 6D).

## Discussion

Temporal prediction has been shown to capture the spatial and temporal tuning characteristics of V1 simple [10] and complex cells [11]. However, these prior modeling studies used feedforward ANNs that omitted recurrence, Dale’s law and all-or-nothing spiking. We explored whether a recurrently-connected, temporal-prediction spiking model incorporating Dale’s law would better replicate the functional properties of V1 neurons, particularly the differences known to exist between excitatory and inhibitory neuron subclasses. Our model captured simple and complex cell-like tuning and V1-like spike statistics to natural stimuli. Notably, the excitatory and inhibitory units in the model exhibited several properties that match their biological counterparts in V1. The much greater explanatory power of the recurrent, spiking model provides strong evidence that temporal prediction may be critical to the function of V1 and potentially other sensory brain areas.

A major goal of computational neuroscience is to develop models that help to interpret the vast amount of experimental data obtained from studies of regions such as V1. This requires creating biologically-realistic models with a functional objective that stipulates how neurons transform visual input [57]. Some previous studies have developed spiking models of V1 with distinct subpopulations of excitatory and inhibitory units. These models have demonstrated V1-like spike statistics and motion tuning [4, 58–61]. However, their parameters were hard-coded to produce these phenomena, rather than being trained to achieve a particular goal like temporal prediction. Consequently, they have provided limited insight into the functional principles underlying the transformation of visual signals in the cortex.

Other studies have shown how specific functional objectives can account for simple cell-like and in some cases complex cell-like properties. These objectives include efficient [62, 63] and sparse coding [64–66], which assume that V1 efficiently encodes sensory input under certain metabolic constraints; independent component analysis [67], which stipulates that V1 reduces input redundancies by encoding independent features within sensory stimuli [68, 69]; predictive coding, which also posits that neurons remove statistical redundancies but in this case by signalling unpredictable features within sensory input [70–73]; temporal coherence [74], slow subspace analysis [75] and slow feature analysis [76, 77], which hypothesize that V1 encodes slowly-varying features within the stimulus; and the information bottleneck hypothesis [78], which suggests that V1 encodes sensory features with maximum mutual information about the future, whilst minimizing information about the past [79]. However, these studies all used non-spiking models and did not include Dale’s law. An exception are sparse coding models that have been extended to be spiking [30, 80, 81] and to include Dale’s law [81]. Although these spiking sparse coding models successfully capture V1 spatial RF properties, they ignore their temporal tuning since the models were trained on images. Temporal prediction has also previously been explored in a spiking form [82], but this study focused on the retina and did not explore Dale’s law.

Whilst the inclusion of spikes and Dale’s law increased the biological realism of our model over prior studies, there remains scope to include additional cortical circuit motifs. The cortex is a 6-layered structure, with layer 4 being the main target for sensory inputs from the thalamus [83, 84]. We currently envisage the units of our model comprising those over all layers. Sub-dividing our model units into a similarly multilayered structure would allow us to examine further phenomena, like the layer-specific tuning characteristics found in V1 [26]. Additionally, our model could be extended to operate over an entire visual scene instead of patches, by jointly training multiple versions of the model at different spatial locations within the visual frame. Incorporating cross-spatial connectivity may segregate the tuning statistics in this model to capture the canonical columnar organization of V1 [2], where neurons with similar orientation preference cluster together [85, 86], or the lateral connectivity statistics, where different neurons are more likely to be connected if they share similar tuning preferences [87, 88]. The optimization of cortex for temporal prediction, at least for the non-hard-wired aspects of cortical tuning, could be guided by the error-sensitive neurons found in cortex [9], or possibly by hypothesized error-sensitive dendrites [89]. These error-sensitive components could compare long (such as from layer 2/3) and short (such as from layers 4 or 5/6) latency neural activity [90] to provide a temporal-prediction error signal [11, 91]. Modeling the learning using more biologically-based local learning mechanisms [92] and structures that are congruent with these phenomena would be another valuable future avenue.

Although our model exhibits V1-like firing dynamics, modelling the units using the adaptive leaky integrate-and-fire model would better match the adaptive spiking dynamics of V1 neurons [15], for example, the decrease in firing observed in response to a constant input stimulus [93]. This would also allow differences in adaptation between excitatory and inhibitory V1 neurons [94, 95] to be explored. In our model, we assumed the effect of input current from one unit to another to be instantaneous. However, synapses conduct current with a temporally-decaying profile, where different types of synapses are governed by a fast or slow decay [96]. Extending our model to include learnable synaptic-like temporal profiles may capture these synaptic phenomena. This would ideally require training our model at a finer temporal resolution than ∼ 8.33ms per time step, which remains a challenge for spiking networks due to speed and memory constraints (although see [97, 98]).

Our temporal prediction model makes several biological predictions. First, based on our finding of a negative exponential relationship between the highest spatial RF power and the membrane time constant across the model units, V1 neurons that are more sensitive to visual input may operate at faster timescales. Second, our findings have implications for calculating the exact energy demands of neurons, which remains an ongoing area of research [99–102]. Specifically, the model predicts that inhibitory V1 neurons may consume less energy per spike than their excitatory counterparts, as we found this assumption to be important for obtaining our results. Third, our results are relevant to the long-standing question of the extent to which visual processing is hierarchical [103, 104] - transforming through sequential layers - or shallow – emerging through recurrent processes. Our model is consistent with the shallow brain hypothesis [105], as it showcases how a single layer of recurrently-connected spiking units captures non-linear phenomena like complex cell-like responses, which have otherwise been argued to emerge from hierarchical processing [106]. This may inspire new computer vision models that favor recurrence over standard hierarchical architectures [107].

In conclusion, we have shown how efficient temporal prediction in a spiking model accounts for simple and complex cell-like tuning, V1-like spike statistics and, notably, key differences between excitatory and inhibitory V1 neurons. Our findings contribute to the growing evidence that visual cortex supports prediction of the sensory future [10, 11]. Although predictive processing is increasingly widely recognized [9], it is perhaps surprising that this single computational principle accounts for several key differences between excitatory and inhibitory V1 neurons. Perhaps many more of the brain’s perplexing phenomena may be explained by the principle of temporal prediction.

## Methods

### Spiking V1 model

#### Training data

We used a publicly available dataset of natural movie recordings to train and test our model [108]. This dataset consists of 39 clips, of 9.6 second duration, of everyday objects and scenes (*e*.*g*. a dog walking, a fish swimming, or a human’s view walking through nature), recorded using different camera movements (*e*.*g*. moving forwards or sideways, panning or still). We used 31 clips for training and 8 clips for testing. All clips are grayscale, with a temporal resolution of 120Hz and a spatial resolution of 140 *×* 240px. To emulate the transformation of the retina and thalamus, we bandpass filtered all clips, as done in other studies [10, 109]. We also normalized the training and test data by subtracting the pixel mean and dividing by the pixel standard deviation of the training data, and finally clipped the absolute pixel values to be no larger than 3.5 standard deviations. Training was performed using clips consisting of 20 *×* 20px spatial patches of 350ms duration (*i*.*e*. 42 frames). The spatial position of each patch were randomly sampled on each training batch. Clips were also randomly flipped around the vertical axis to artificially increase the training data [110].

#### Spiking neurons

We modeled the V1 cortical neurons as a population of *N* = 600 recurrently-connected excitatory and inhibitory spiking units. The input current *I*_*i*_[*t*] ∈ ℝ to unit *i* at time step *t* is derived from the bandpass-filtered input movie stimulus *x* ∈ ℝ^*T ×H×W*^ (of *T* = 42 frames, and spatial height *H* = 20px and spatial width *W* = 20px); model output spikes *S*[*t* − 1] ∈ ℝ^*N*^ from the previous time step; and a bias term 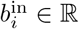.

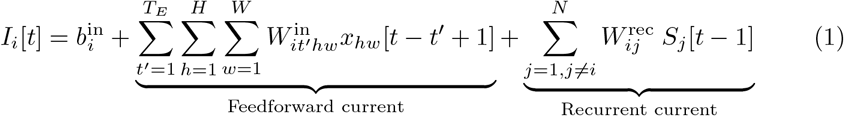

The feedforward connectivity 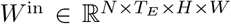 (of *T*_*E*_ = 15 temporal frame span) linearly maps the input movie stimulus into a feedforward current contribution (capturing the transformation of the retina and thalamus); the recurrent connectivity *W* ^rec^ ∈ ℝ^*N×N*^ maps previous output spikes into a recurrent current contribution. We implemented all spiking units using the normalized and discretized leaky integrate-and-fire (LIF) model, which evolves the model membrane potential *V*_*i*_[*t*] ∈ ℝ of unit *i* at time step *t* using the following difference equation:

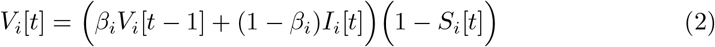

During each time step, the membrane potential undergoes a decay governed by a factor *β*_*i*_. We let each neuron learn its decay factor, and bounded values into the range (0, 1) to enforce correct LIF dynamics [111]. *I*_*i*_[*t*] was replaced with *Ĩ*_*i*_[*t*] when synaptic noise was included. The membrane potential resets to zero in the event of a spike occurring in the preceding time step, which happens when the membrane potential reaches the firing threshold (equal to one).

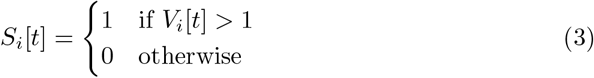

Finally, at every time step *t*, a prediction of the future spatial movie frame *ŷ*[*t*] ∈ ℝ^*H×W*^ was generated (with *ŷ*_*hw*_[*t*] ∈ ℝ denoting the predicted pixel value). This was achieved by linearly mapping a span of *T*_*D*_ = 2 temporal frames of proceeding spike activity using projection weights 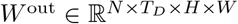, plus a bias *b*^out^ ∈ ℝ.

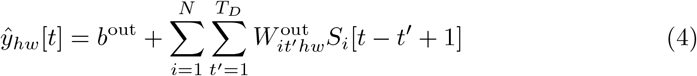

#### V1 latencies

V1 neurons have a response latency of at least ∼ 30 − 40ms in mice and monkeys [17–20], and these latencies can vary substantially between neurons [112]. We modeled the transduction and transmission latency to V1, by zero-masking the first 5 spatial frames (*i*.*e*. 42ms) within the feedforward connectivity tensor *W* ^in^. This length is the temporal prediction span.

#### Neural noise

Cortical V1 neurons do not respond with identical spike trains to repeated visual stimulus presentations. One of the reasons for this is neural noise. We include two neural noise sources within our model. First, we modeled the noisy photoreceptors in the retina [113], by adding Gaussian noise to the bandpass-filtered movie clips *ϵ*_*p*_ ∼ *𝒩* (0, 0.2^2^). Second, we modeled the synaptic noise of cortical V1 neurons [114], by multiplicatively perturbing the input current *I*_*i*_[*t*] of every unit *i* at time step *t* using Gaussian sampled noise *ϵ*_*g*_ ∼ *𝒩* (0, 0.6^2^).

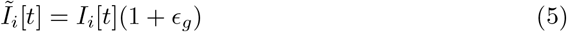

#### Dale’s law: Enforcing units to be inhibitory or excitatory

Approximately 15 − 20% of cortical V1 neurons are inhibitory in mice, cats, and monkeys [5–8]. We assigned 15% of the model unit population to be inhibitory, resulting in *N*_*I*_ = 90 inhibitory units and *N*_*E*_ = 510 excitatory units. We enforced a neuron to be excitatory (or inhibitory), by taking the absolute value of every weight in the unconstrained recurrent weight matrix 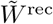, and appropriately negating connection weights from unit *j* to *i*. All outwards connection weights from unit *j* are either positive (*i*.*e*. excitatory) or negative (*i*.*e*. inhibitory).

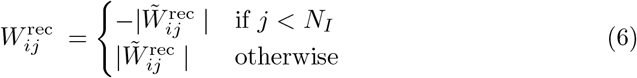

#### Weight initialization

The initial weights for feedforward (*W* ^in^), recurrent (*W* ^rec^), and projection (*W* ^out^ - *i*.*e*. the readout weights) connections were randomly initialized from a uniform distribution *U* (−*k, k*), where *k* was determined as 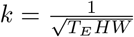 for the feedforward weights, 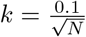 for the recurrent weights, and 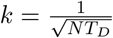 for the projection weights. Additionally, all initial membrane time constants were uniformly initialized to 20ms (*i*.*e. β*_*i*_ ≈ 0.81 for all units). Lastly, the input bias terms were initialized to 0.2 and the output bias term to 0.

#### Loss function

We trained the model by minimizing loss ℒ_total_, comprised of prediction loss ℒ_normative_ and metabolic loss ℒ_metabolic_, with respect to *W* ^in^, *W* ^rec^, *W* ^out^, *b*^in^, *b*^out^, and *β*. The prediction loss measures the similarity between the predicted and target future movie frames, and the metabolic loss measures the approximated physiological energy use of the network. We weighted this loss by hyperparameter *λ* = 10^−2.75^, which we selected to produce the most V1-like receptive fields.

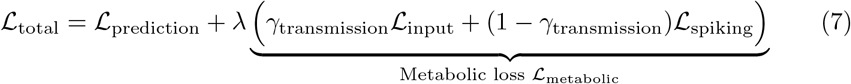

We computed the prediction loss ℒ_prediction_ as the mean squared error (MSE) between all predicted spatial movie frames *ŷ* and target spatial movie frames *y* ∈ ℝ^*H×W*^. The calculation was performed by taking the average over all batch samples *B* (omitted here for brevity), simulation steps *T* (starting from simulation step *t*_0_ = 5 to allow unit membrane potentials to depolarize sufficiently, *i*.*e*. to enable network warm-up), and spatial dimensions *H* (height) and *W* (width), both which were cropped by *c* pixels on all sides.

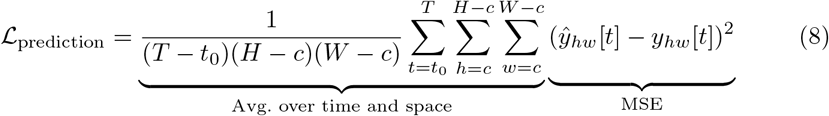

We defined our metabolic-like loss ℒ_metabolic_ as the average synaptic-like transmission over all the model weights, where synaptic transmission has been noted to require significant energy within cerebral cortex [102]. We defined the energy cost of the synaptic-like transmission of a single model weight as its absolute value, multiplied by the absolute value of every input value, summed over all time steps. We weighted the recurrent transmission (ℒ_spiking_) relative to the input transmission (ℒ_input_) using hyperparameter *γ*_transmission_ = 0.3. We also assumed the metabolic-like cost of the excitatory and inhibitory units to differ, with the inhibitory units consuming less energy than excitatory units, captured by hyperparameter *γ*_type_ = 0.1.

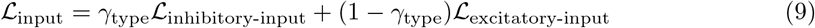

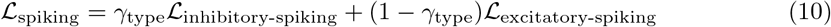

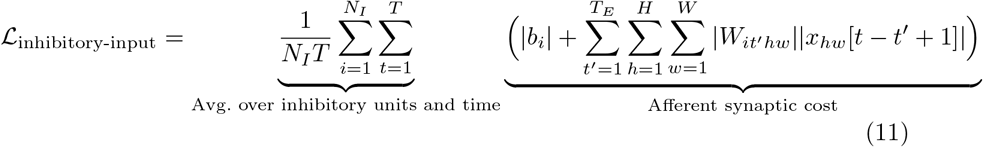

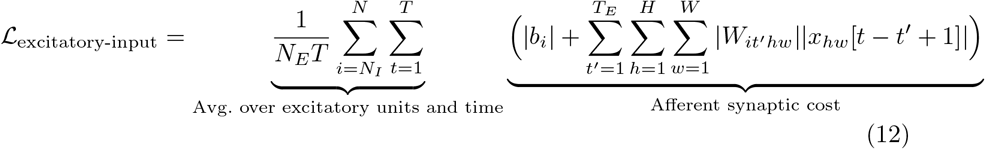

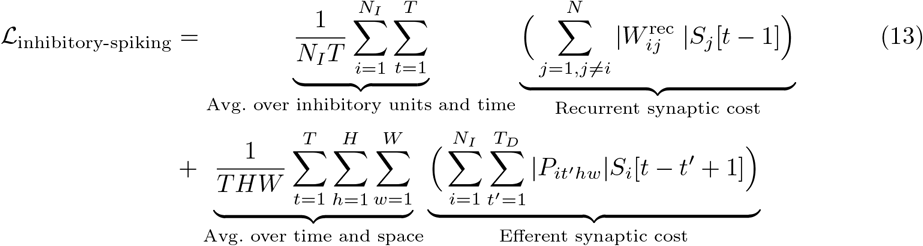

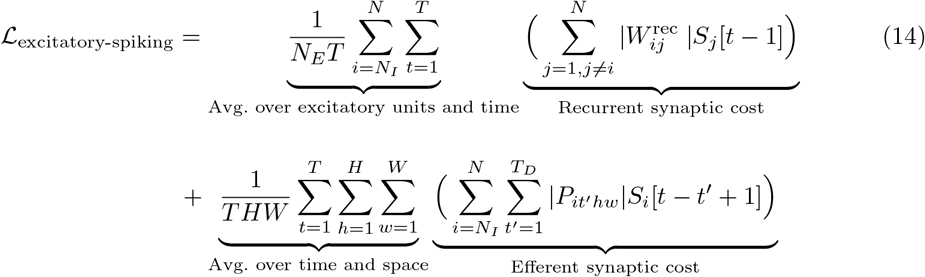

#### Surrogate gradient descent

We trained the spiking models using a variation of the backpropagation algorithm [22] called surrogate gradient descent [23]. Here, to enable smooth gradient flow during training, a well-behaved function was employed as a replacement for the gradient of the non-differentiable Heaviside step function. We adopted the fast sigmoid function, which has been shown to perform well in practice [115, 116].

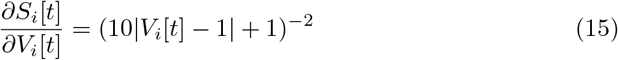

We conducted all training using the Adam optimizer [117], employing its default parameters. The training spanned 1200 epochs with an initial learning rate set to 10^−4^, which was subsequently reduced by a factor of 0.2 at epoch count 200, 600, and 800. We performed training using a batch size of 1024 and saved model weights whenever a new minimum training loss was achieved.

#### Spike analysis

For all the spike analysis, we used the model output spike trains resulting from 8000 different retina-filtered movie patches taken from the Star Wars movies. We used these natural movies in our spiking analyses to be comparable to [4], where they calculated V1 spiking statistics in monkeys watching segments of the Star Wars movies. For additional comparability with the analysis of [4], we calculated all spike statistics using a time window of 2s with a step size of 0.2s.

#### Coefficient of variation

We quantified the variability of the spike pattern in each spike train by calculating the coefficient of variation of the interspike interval CV(ISI), which is defined as the ratio between the standard deviation and the mean of the interspike intervals [15]. A *CV* (*ISI*) *<* 1 corresponds to a more regular firing pattern than a *CV* (*ISI*) *>* 1, and a Poisson process has *CV* (*ISI*) = 1. As done in [4], we only calculated CV(ISI) values for a given spike train if it had at least three spikes.

#### Neuron synchronization

We quantified the correlation between different unit spike trains by calculating the mean cross-correlogram between the spike trains of randomly-sampled unit pairs. This measures the average Pearson correlation coefficient 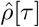 between the spike trains *s*_1_ and *s*_2_ of different units for varying temporal lag *τ*, defined as:

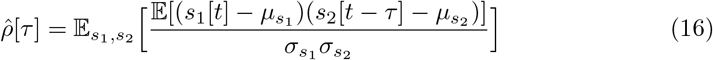

We sampled 400 random unit pairs for this calculation. Furthermore, we binned the spike trains with a bin size of 25ms before calculating the correlations, in order for our analyses to be comparable to [4].

### Receptive field analysis

#### Spike-triggered average

We estimated the spatiotemporal receptive field of every unit using a spike-triggered average [96], using responses to 1000 spatiotemporal white-noise input clips of 100-frame duration. Clips were generated by sampling each pixel from a Gaussian distribution with a standard deviation of *σ* = 10.

#### Fitting Gabors

We fitted a Gabor function [118] to the RF of every unit to quantify its tuning properties. The Gabor function has been shown to provide a good approximation for most RF spatial aspects [32, 33] and provides a quantitative measure to compare the RFs of the model with the RFs of V1 neurons. The 2D Gabor function is defined as:

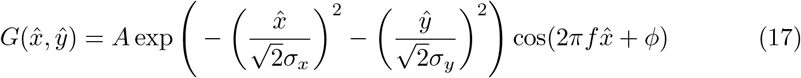

where spatial coordinates 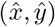 are obtained by translating the center of the RF (*x*_0_, *y*_0_) to (0, 0) and rotating the RF by its orientation *θ*:

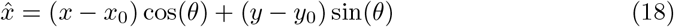

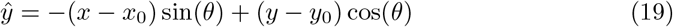

The amplitude of the Gaussian envelope is defined by *A*, whereas parameters *σ*_*x*_ and *σ*_*y*_ define the Gaussian envelope width along the 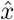 and *ŷ* axes, respectively. Parameters *f* and *ϕ* define the spatial frequency (in cycles/pixel) and phase of the sinusoidal grating along 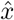. We optimized the Gabor parameters for each model unit RF by minimizing the mean squared error between the corresponding Gabor function and its respective model RF. To avoid local minima during fitting, we performed each Gabor fit in the spectral domain, by transforming the model RFs and Gabors using a 2D Fourier transform [10, 32]. We then used the resulting parameters to continue the fitting procedure in the spatial domain. We excluded units in Figure 3C which had a poor fit (correlation coefficient *<* 0.6, 381 units), whose fit was based on very few pixel values (Gaussian envelopes *σ*_*x*_ *<* 0.5 or *σ*_*y*_ *<* 0.5, 382 units), and whose center position (*x*_0_, *y*_0_) of the Gabor was outside the RF (9 units). This resulted in 141 out of 519 active units with a sufficient fit, having a median pixel-wise correlation coefficient of 0.83 (81 out of the 600 model units were deemed inactive, see Motion tuning section).

#### Spatial receptive field properties

##### Power

The power of a RF is defined as the mean of the squared values of the RF over a span of time (as in Figure 3D) or at a point in time (as in Figure 3E) [10].

##### Structure

The structure of a RF is characterized by two variables *n*_*x*_ = *σ*_*x*_*f* and *n*_*y*_ = *σ*_*y*_*f*, which give a measure of the number of oscillations of its sinusoidal component and the length of the bars in the RF, respectively [33]. Blob-like RFs lie close to the origin in the *n*_*x*_ by *n*_*y*_ plane, the number of bars increases along the *n*_*x*_ axis, and the RF bar-length increases along the *n*_*y*_ axis.

#### Estimating space-time separability

Spatiotemporal RFs are either space-time separable or inseparable, with space-time separable RFs tuned to phenomena such as bars flashing or moving regardless of direction, and space-time inseparable RFs tuned to direction of movement. Space-time separable spatiotemporal RFs can be represented as a product of a spatial and temporal function while space-time inseparable spatiotemporal RFs cannot [96]. We classified the space-time separability of model unit RFs using the singular values of a singular value decomposition (SVD), as done in [10]. We flattened the spatial dimensions and removed the transmission latency time steps for each model unit RF, and took the SVD of the resulting matrix. If the ratio between the second and first singular value was *<* 0.5, the RF was deemed space-time separable; otherwise, the RF was deemed space-time inseparable.

#### 2D spatiotemporal receptive fields

The 2D spatiotemporal RF illustrates the temporal evolution of the spatiotemporal RF [35]. This view was constructed by translating and rotating every spatial RF by the position and rotation that centers the spatial RF of the largest power, placing all oriented bars parallel to the y-axis. Next, all spatial RFs were flattened along the y-axis and concatenated, resulting in a 2D spatiotemporal RF. This new view can qualitatively inform us about the temporal phase (through polarity changes in the flattened spatial structure) and separability (through shifts in the flattened spatial structure) of a spatiotemporal RF.

### Motion tuning

#### Virtual physiology

We constructed model unit responses to drifting full-field sinusoidal gratings, varying in orientation (from 0° to 360° in 5° increments), spatial frequency (ten values evenly spaced from 0.01 to 0.2 cycles per pixel), and temporal frequency (1, 2, 4, and 8Hz), each with an amplitude of one. Each grating clip, lasting 3 seconds, was presented to the model four times. We calculated the firing rate of each model unit by averaging responses over clip repeats and convolving with a Gaussian kernel (*σ* = 72ms) [96]. For each model unit, we computed its mean firing rate over each grating clip, and constructed a 3D mean-firing tuning space (over orientation, spatial, and temporal frequency), with the optimal grating (defined by its orientation, spatial and temporal frequency) eliciting the largest mean firing rate. We only included responsive units in the tuning analysis of Figure 5 (519 out of 600 units) that had an optimal response no less than 10% of the mean optimal response over all model units.

#### Modulation ratio

We measured the modulation ratio *F*_1_*/F*_0_ of each model unit by taking the Fast Fourier Transform (FFT) of each unit’s response to its optimal grating and calculating the ratio between the amplitude of the first harmonic (*F*_1_) and DC component (*F*_0_). A unit with a modulation ratio below one is considered complex cell-like, whereas a unit with a modulation above one is considered simple cell-like [26, 40, 41].

#### Orientation and direction selectivity

We calculated the orientation selectivity index (OSI) and direction selectivity index (DSI) for each unit, which quantify a unit’s preference for stimulus orientation and direction of motion, respectively [26, 44]:

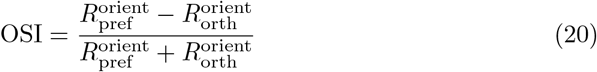

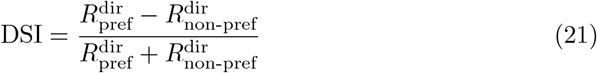

Both metrics are calculated using the mean unit responses to gratings at a unit’s preferred spatial and temporal frequency. 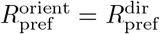 is a unit’s mean response to a stimulus in its preferred orientation and direction, and 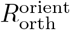 and 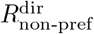 are the mean responses in the orthogonal orientation and opposite direction to the preferred direction, respectively. A value close to zero corresponds to less selective responses, and a value closer to one corresponds to greater selectivity for both measures.

### Physiology analysis

#### Calculating membrane time constants

The membrane time constant *τ* of every model unit is obtained from the decay factor *β* in the LIF Equation using the following relation:

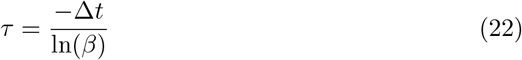

**EI balance**

Using the held-out test dataset of natural movie stimuli, we calculated the net-excitatory (Ex_*i*_) and net-inhibitory (In_*i*_) input current to each model unit *i* at time step *t* as:

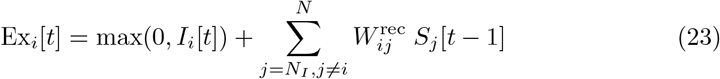

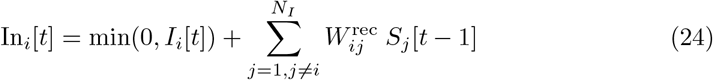

The net-excitatory and net-inhibitory currents were derived from the excitatory and inhibitory units, respectively, with the feedforward current *I*_*i*_[*t*] being classed as either excitatory or inhibitory at a given time step, depending on its sign value. From this, we calculated the global EI balance of a model unit as:

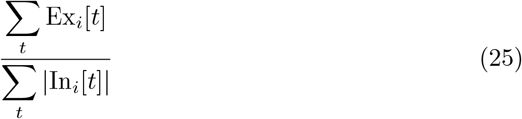

with the excitatory and inhibitory current time series for each input clip concatenated over time.

We calculated the precise EI balance of every model unit by first convolving each current time series using a Gaussian kernel (*σ* = 72ms) [96] and then calculating the resulting correlation between the smoothed (negative) inhibitory and (positive) excitatory current time series (with both time series concatenated over time for each input clip). We smoothed the current time series to emulate synaptic conductance [15], finding the non-smoothed excitatory and inhibitory current time series of units to otherwise be uncorrelated (median CC of 0.04).

## Acknowledgments

Luke Taylor was supported by the Clarendon Fund of the University of Oxford. Andrew King and Nicol Harper were supported by the Wellcome Trust (WT108369/Z/2015/Z). Friedemann Zenke was supported by Swiss National Science Foundation (grant no. PCEFP3 202981) and the Novartis Research Foundation.

## Data availability

The natural movie training data can be downloaded from https://figshare.com/articles/dataset/Natural_movies/24265498 and movie test data used in the spiking analyses can be downloaded at https://github.com/webstorms/V1Model. The untrained and pre-trained spiking models can be found at https://github.com/webstorms/V1Model. Experimental data in Fig. 5 was extracted using an online tool available at https://apps.automeris.io/wpd/.

## Code availability

The code of the spiking V1 model and the code for reproducing the results can be found at https://github.com/webstorms/V1Model.

## Notes

https://github.com/webstorms/V1Model

### Competing Interest Statement

The authors have declared no competing interest.

